# A standardized non-visual behavioral event is broadcasted homogeneously across cortical visual areas without modulating visual responses

**DOI:** 10.1101/2020.12.15.422967

**Authors:** Mahdi Ramadan, Eric Kenji Lee, Saskia de Vries, Shiella Caldejon, India Kato, Kate Roll, Fiona Griffin, Thuyanh V. Nguyen, Josh Larkin, Paul Rhoads, Kyla Mace, Ali Kriedberg, Robert Howard, Nathan Berbesque, Jérôme Lecoq

**Author notes:** **Correspondence and requests for materials** should be addressed to M.R. or J.L. **Competing interests** The authors declare no competing interests.

## Abstract

Multiple recent studies have shown that motor activity greatly impacts the activity of primary sensory areas like V1. Yet, the role of this motor related activity in sensory processing is still unclear.

Here we dissect how these behavior signals are broadcast to different layers and areas of the visual cortex. To do so, we leveraged a standardized and spontaneous behavioral fidget event in passively viewing mice. Importantly, this behavior event had no relevance to any ongoing task allowing us to compare its neuronal correlate with visually relevant behavior like running.

A large two-photon Ca^2+^ imaging database of neuronal responses uncovered four neural response types during fidgets that were surprisingly consistent in their proportion and response patterns across all visual areas and layers of the visual cortex. Indeed, the layer and area identity could not be decoded above chance level based only on neuronal recordings. In contrast to running behavior, fidget evoked neural responses were independent to visual processing.

The broad availability of visually orthogonal standardized behavior signals could be a key component in how the cortex selects, learns and binds local sensory information with motor outputs. Contrary to relevant motor outputs, irrelevant motor signals would use a separate neural subspaces.

**Significance Statement:** Recent studies have shown contextual and behavioral variables to dominate brain-wide activity, but yet it is unknown how this information is broadcast across cortical layers and areas. Using a large two-photon dataset collected in mice passively viewing a battery of visual stimuli, we characterized the neuronal response of neurons of the visual cortex to a standardized fidget behavior. We found that as much as 47% of excitatory neurons show significant co-activity with fidgets. Throughout all areas and layers we recorded from, those responses were distributed across surprisingly consistent three neural response types. Further analysis showed no interaction between fidget neural events and cells’ visual stimulus responses, contrasting with feedback neural signals induced by running. These contrasting response distribution patterns suggest that behavioral neuronal correlates are broadly available but will modulate sensory responses depending on their relevance to local sensory inputs.

## Introduction

Traditionally, the sensory cortex has been modelled as a feed-forward structure where low level information, e.g. pixel-wise visual inputs, are integrated with global behavior signals like motor output downstream of the visual cortex^29^. Many principles of deep learning were inspired by this view and fueled the modern rise of artificial networks. Indeed, initial reports of visual response modulation in V1 show weak modulation by behavior in monkeys^1^. This result was experimentally challenged in mice with the discovery of strong running modulation in V1^2^ as well as across multiple sensory areas^3–6^. In addition, it was demonstrated that non-visual events are not mere modulators but can also directly evoke neuronal activity in V1^7^. In fact, contextual and behavioral variables have recently been shown to largely dominate brain-wide activity^8,9^. This result brings into question the role of these events. If the brain broadcasts behavior relevant variables like motor outputs, this should allow each brain area to integrate this information into its computation^5^. As a result, understanding the micro-circuit computation occurring across all cortical layers and cell types in this context requires a detailed physiological characterization of the neuronal correlate of motor outputs.

One approach to tackle this challenge is to monitor all potential behavior events and characterize all associated neuronal correlates^1^. This is challenging since motor outputs, contrary to sensory stimuli, are highly variable from trial to trial and hard to standardize and control. In addition, behavioral events like running are correlated with a complex symphony of sensory-motor events, tangling together feedforward visual and visually-relevant feedback signals. Running behaviors are also inherently associated with visual motion, triggering visual predictions mismatch in head-fixed mice^7^. Thus, to extract principles on the neuronal correlates of behavioral events, it is helpful to investigate a non-visually relevant behavioral event. Here our goal is to characterize how a standardized behavior output differently affects all areas, layers and cell types of the cortex in order to provide foundational knowledge for modelling cortical computation. To achieve this goal, we leveraged the natural occurrence of fidgets in experimental mice.

Fidgets are stereotypical behavioral responses that are potentially part of a stressful state^10^. They manifest as a fast, spontaneous, startle response accompanied with stereotypical body chest movements. We used fidgets detected in behavioral videos to quantify and compare the neuronal correlates of standardized behavior events across all layers and most areas of the mouse visual cortex. To this end, we leveraged a large survey of neuronal responses recorded with *in vivo* two photon calcium imaging in the mouse visual cortex^4^. While the brain-wide impact of behavioral events is now established, our analysis revealed that neurons in three different cortical layers and four visual areas have homogenous post-fidget neural responses. Fidget response profiles were stereotypical and equally distributed among 4 response types. In addition, in contrast to running behaviors, visually evoked neuronal responses showed no interaction with fidget neural events.

## Results

### Fidget as a standardized behavior output

We first sought to characterize the range of behavioral events that mice displayed under head fixation. While we recorded neuronal activity *in vivo* using two photon imaging, mice were free to run on a rotating disc while a camera captured mouse body posture **(Fig. 1a-c)**. We observed a variety of behaviors such as whisking, grooming, mastication, flailing (uncoordinated movement), walking, running, and a startle behavior we denote as a “fidget”. Fidgets manifested as a combination of abdominal flexion (causing the abdomen to be raised above the rotating disk) and an upward force generated from the lower limbs causing lower trunk curvature and contraction **(Fig. 1d)**. Fidgets were qualitatively stereotyped across mice in their duration, pattern of movement, and motor response magnitude. Following this observation, we sought to develop a computer vision model to automatically identify fidget events from hours of mice behavioral videos. Six human annotators first established a training data set (20 mice, 10,000 fidgets manually annotated, see **Methods**). We computed the Histograms of Oriented Gradients (HOGs) for each video frame and concatenated a feature vector from a one second section of frames (30 frames) **(Fig. 1e)**. HOGs are transformation invariant visual features extracted using edge-detection-like computation. HOG features are largely invariant to variation in lighting conditions and image transformations such as translation, rotation, scale. This allows us to carry out robust behavioral feature detection as others have firmly established^28^. Another advantage of using HOG vectors is the biologically inspired emphasis on interpretable edge detection computations and has been found to be superior to Eigenfeature based face recognition models^27^.

**Fig. 1.**
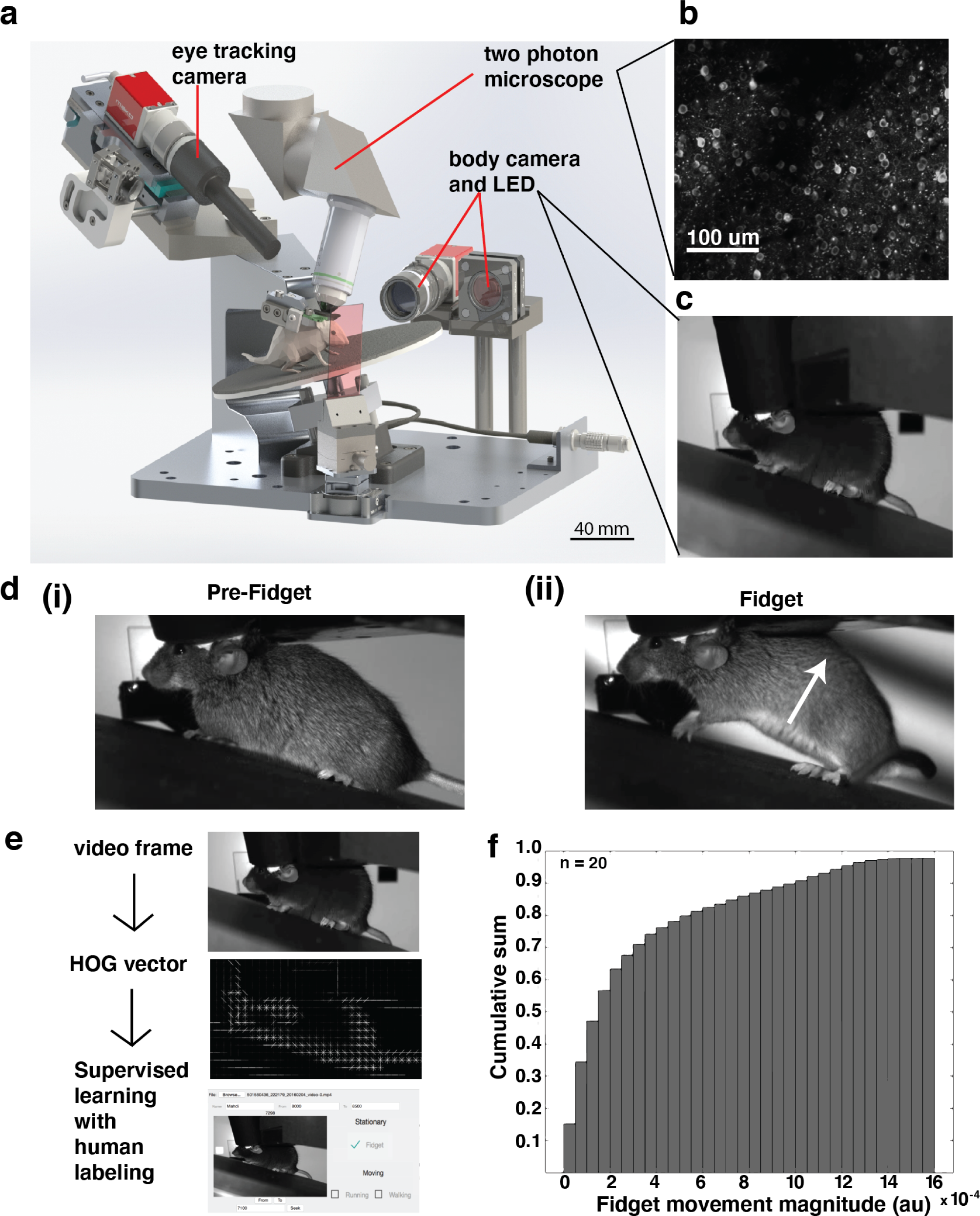
Detection of behavioral fidget during two photon imaging in freely-viewing mice. **(a)** Computer design of our apparatus to monitor the behavior of mice during head-fixation and two-photon imaging. **(b)** Example two photon imaging field of view (400 μm x 400 μm) showcasing neurons labeled with Gcamp6f. **(c)** Example video frame of mouse captured by the body camera at 30Hz. **(d)** A pair of video frames showing the progression of a prototype fidget behavior in time. **(i)** First, the mouse is stationary **(ii)** Then, during the initiation of the startle response, the mouse stereotypically pushes its body up using its bottom paws while arching the back and contracting the abdomen. **(e)** Computational strategy for detecting fidgets automatically. **(f)** Cumulative probability sum of labelled fidget moment magnitude, showcasing the consistency of the stereotyped behavior across experiments and mice. Fidget magnitude is calculated as the sum of 2D optical flow vectors throughout the fidget duration (see **Methods** for optical flow calculation).

Using this feature, we trained a Support Vector Machine (SVM) classifier in a supervised manner using the human-annotated labels. Our final trained model had a recall performance of **74% +/- 4.2 (mean +/- std, n=7)** and a precision performance of **78% +/- 5.3 (mean +/- std, n=7**) for the seven one-hour long experiments held out as a validation test-set. Our trained classifier was as accurate at identifying fidgets and other mouse behaviors as human annotators. Indeed, seven pairs of annotators analyzed the same videos and their annotations were compared head-to-head. Each video was drawn from a seven video validation test-set. Head-to-head human vs. human performance recall for the seven videos was **73% +/- 5.9 (mean +/- std, n=7)** and had a performance precision of **74% +/- 7.2 (mean +/- std, n=7)**; this was within the range of the model’s performance. Having established a robust computer vision model, we automatically annotated 144 one-hour experiments total (recall p = 0.51, performance p = 0.26).

To quantify the standardization of fidget events across mice, we integrated the optical flow magnitude of the fidget motor response (see **Methods**) over the duration of the fidget. 80% of all fidget events from 20 one-hour experiments and across 20 unique mice fell within 30% of the maximum magnitude; this consistency reaffirmed the stereotypy of fidget events **(Fig. 1f)**.

### Occurrence of fidget across mice and visual stimuli

We next sought to establish whether the occurrence of fidgets could relate to our visual stimuli. Fidget behavior has been associated with stress and surprise responses in mice^10,14-15^. As described in a previous publication^4^, mice passively viewed a range of both artificial (drifting and static gratings, locally sparse noise) and natural visual stimuli (natural scenes and natural movies), organized into three different recording session (sessions A,B and C) **(Fig. 2a)**. We hypothesized that artificial visual stimuli (e.g. drifting gratings) induce a more stressful or surprising context than natural stimuli that are more ethologically familiar to the mouse (e.g. natural movies). In particular, the moving drifting gratings, through its perceived motion, evoke an innate avoidance response^3^. In line with our prediction, the average normalized fidget rate was significantly higher during drifting gratings **(Fig. 2a**, p = 0.023, two tailed t-test, n = 60) than other stimuli **(Fig. 2b)**.

**Fig. 2.**
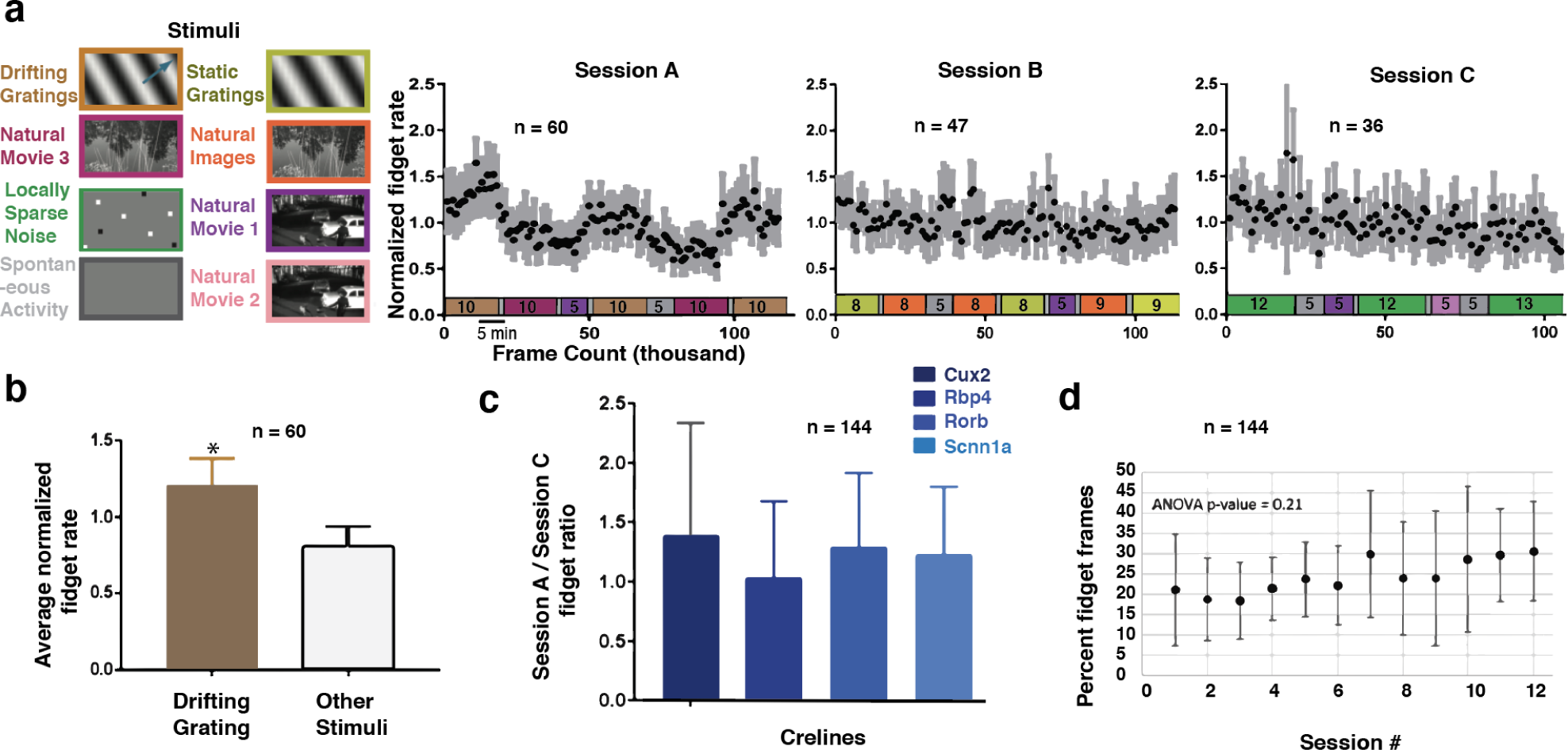
Fidget rate is correlated with visual stimulus type, but independent of a mouse driver-line or session number. **(a)** (left) Standardized experimental design of sensory visual stimuli. Six blocks of different stimuli were presented to mice and were distributed into three separate protocols identified as Session A, Session B, and Session C (right). Normalized fidget rate (black dots) for all three session types across all mice. Grey bars indicate 95% mean confidence intervals. Color coded stimulus protocol with indicated durations (minutes) are aligned with the time axis for all three session types. **(b)** Average normalized fidget rate across all mice during the presentation of drifting grating visual stimulus in comparison to all other stimuli (p-value = 0.023). **(c)** Comparison of the fidget rate ratio between different session types across cre-lines. No significant effect was found (ANOVA, p-value = 0.14). **(d)** average percent of video frames labeled as fidget vs. the number of experimental sessions a mouse has been exposed to, no significant learning effect found (ANOVA, p-value = 0.21).

We found that fidget rate was highly variable across mice (see **Supplementary Fig. 1)**, raising the possibility that various mouse Cre-lines have different stress sensitivities, thus accounting for the fidget rate variance. The absolute fidget rate did not significantly differ between mice from different Cre-lines (ANOVA p-value = 0.14, n = 144). This result did not exclude that mice could be more sensitive to individual stimuli. To account for variability across individual mice, we normalized the change in fidget rate evoked during the session with drifting gratings (session A) by the fidget rate during session C **(Fig. 2c)**. Similarly to the absolute rate, we saw non-significant changes (ANOVA p-value = 0.14, n = 144).

Previous research has shown that fidget behaviors can be learned^10^. Our passive viewing protocol included two weeks of habituation to our visual stimuli (see **Methods**), suggesting we could have reached a more stable state. To check whether the fidget occurrence we see is learned over the course of our two-photon experiments, we quantified the average fidget rate across mice as a function of the number of visual stimulation sessions the mouse has already seen. We found a non-significant change in the fidget rate with an increased number of sessions experienced **(Fig. 2d)**, supporting the claim that we are operating in a stable behavioral regime and that the mice are adequately habituated (ANOVA p-value = 0.21, n = 144).

In summary, we saw no significant difference in the fidget rate between Cre-lines but different visual stimuli evoked different fidget rates. Importantly, there were no learning effects as mice were already habituated to the stimulus. These results allowed us to explore the neuronal correlates of fidgets across layers and areas of the visual cortex with two-photon calcium imaging.

### Neuronal correlates of fidgets

We analyzed experiments where adult mice (90 +/- 15 days) expressed a genetically encoded calcium sensor (GCaMP6f) under the control of specific Cre-line drivers (Rorb, Cux2, Rbp4 and Scnn1a excitatory lines, see **Methods**). Data was collected from four visual cortical areas (VISal, VISI, VISp, VISpm) and three different cortical layers (Layer II/III, layer IV, and layer V; 175 μm, 275 μm, and 375 μm depth respectively)^2^. In total, we analyzed the activity of 20,253 neurons imaged during 144 one-hour imaging sessions. Visual responses of neurons at the retinotopic center of gaze were recorded in response to drifting gratings, flashed static gratings, locally sparse noise, natural scenes and natural movies displayed on a screen. The analysis of these visually evoked responses was published previously^2^. Here we focused on the neuronal correlates associated with fidgets and their overlap with a subset of visual stimuli.

Many neurons showed a robust and prolonged response after fidget onset, in line with multiple previous studies studying the brain wide effects of motor responses^2,8,16-17^ **(Fig. 3b)**. We first checked that this large neural response was not due to motion artifacts caused by the fidget itself (See **Supplementary Figure 3**). First, these calcium events were prolonged and delayed, a temporal dynamic incompatible with an immediate motion artifact. In addition, many fidgets evoked clear global events across large portions of the field of view with minimal movements of the 2p image (see **Supplementary Videos 1**).

**Fig. 3.**
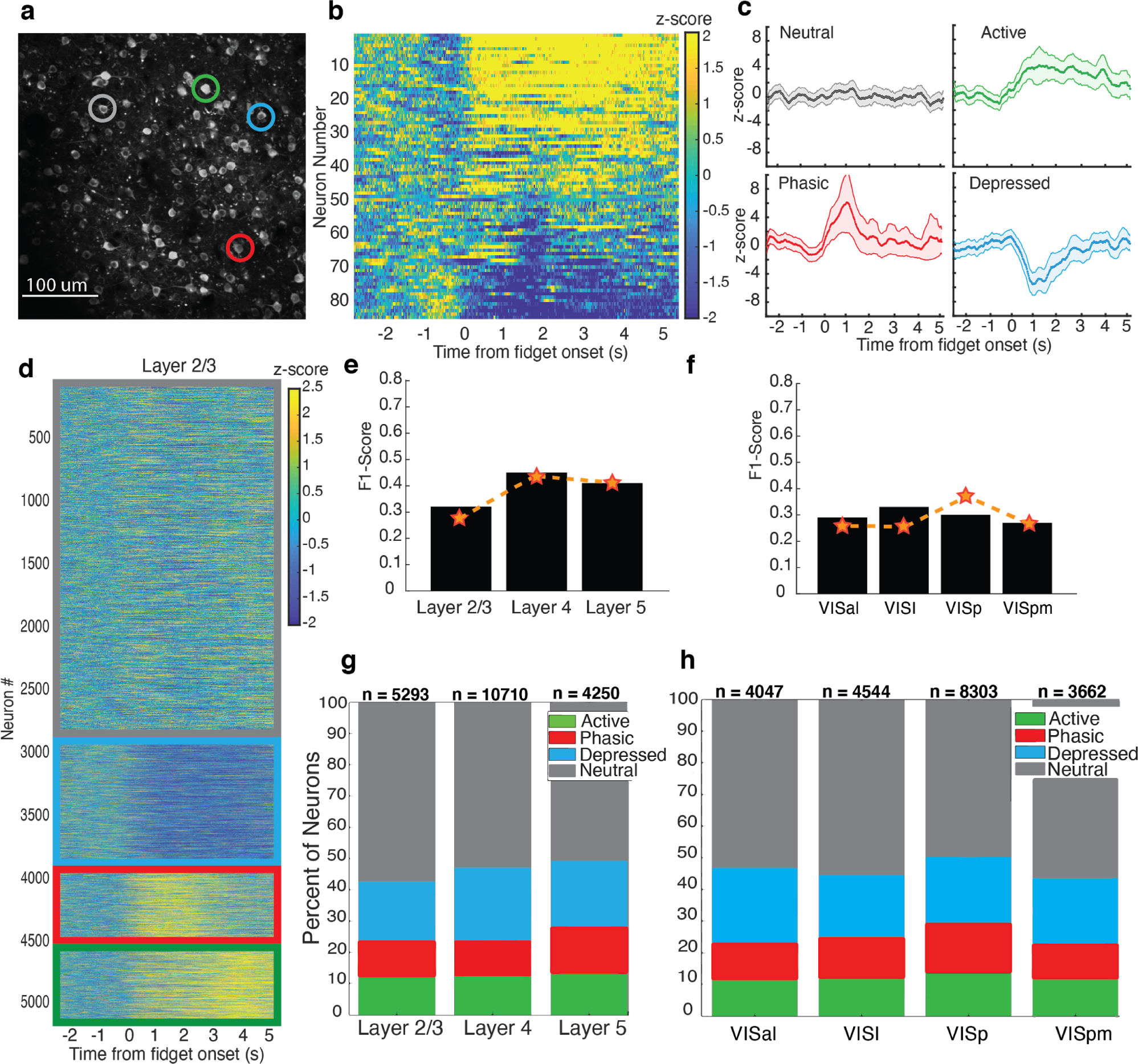
Neuronal fidget response types are distributed equally across visual cortical layers and areas. **(a)** Example two photon imaging field of view (Cux2, 400 μm x 400 μm) showcasing all neurons recorded in one session. Four unique neuronal response types are displayed by the four neurons identified by colored circles. Scale bar = 100 μm. **(b)** The trial averaged z-scored activity of all neurons from one experiment, aligned to the time of fidget initiation (0 seconds). **(c)** The activity profiles of exemplary neurons showcasing the four types of neuronal responses identified using clustering (see **Method**). Top-left in grey: neutral, top-right in green: active, bottom-left in red: phasic, bottom-right in blue: depressed. Colored points indicate the trial averaged z-scored activity, shaded region indicate one standard deviation across trials for each time point. **(d)** All layer 2/3 neurons from all experiments clustered into the four neuronal response types outlined by color coded borders (neutral: grey, depressed: blue, phasic: red, active: green). **(e)** Decoding of neural response type using UMAP feature vector across cortical depths (see **Method**). Results suggest that the neural response types cannot be differentiated across cortical depth (black bars: F1 score based on neural data, orange dashed line: F1 score based on shuffled neural data) **(f)** Decoding of neural response type using UMAP feature vector across cortical areas. Results suggest that the neural response types cannot be differentiated across cortical area (black bars: F1 score based on neural data, orange dashed line: F1 score based on shuffled neural data) **(g)** Percent distribution of neuronal response types per cortical layer for all cortical areas. **(h)** percent distribution of the neuronal response types per cortical area for all cortical layers.

To investigate the structure of these seemingly global neural events, we looked at individual neuron responses across trials. Interestingly, neuronal responses fell into four distinct types based on the direction, magnitude and durations of the response: *Neutral* neurons did not have activity changes from pre-fidget to post-fidget, *Phasic* neurons displayed a transient increase of activity right after fidget initiation, *Active* neurons maintained this increase throughout the post-fidget period, while *Depressed* neurons displayed a decrease in activity post-fidget. These response patterns were stable across trials for each neuron and displayed little deviation from the trial mean. Similar response clusters have been identified previously^18-20^ **(Fig. 3c)**.

### Neuronal fidget responses across layers and areas are uniform

Although it is now established that behaviorally related neural events are evoked in the visual cortex, we wanted to evaluate if these neural response types were localized to a certain layer or area of visual cortex. To quantify how these response types were distributed, we first used a time series k-means clustering algorithm (see **Methods**). Across all 144 experiments, we found a surprisingly large portion of neurons whose activity was impacted by fidgets. 47.2 percent of neurons were classified as active (12 %), phasic (13.9%) or depressed (21.3 %), and 52.8% as neutral. The clustered activity profiles displayed high consistency within each layer and area **(Fig. 3d)**, and the clustering was able to generalize with high accuracy across layers and areas **(Fig. 3e-f)**. Crucially, when conditioned on different layers and areas, the distribution of neural response types was surprisingly consistent, with around 47% of neurons on average being classified as active, phasic or depressed (**Fig. 3g-h**). To verify our result was not impacted by a selection criteria on neurons to be clustered, we applied several different criteria derived from previous publications^23^ to our clustered neural data as a control. As expected, the percent of neurons significantly modulated in the post fidget period spans from ~8% to ~25% of all neurons depending on the strictness of the threshold criteria. Crucially, the distribution of neural response types remained consistent when conditioned on a threshold criteria for different layers and areas (See **Supplementary Figure 4**).

We next investigated whether layers or visual areas were differentially modulated by what could be a behaviorally relevant feedback input. To do so, we projected each 200-dimensional post-fidget neutral response (each time point being a feature) into a 2-dimensional subspace found using UMAP, a non-linear dimensionality reduction technique^26^. If cortical layers or areas differed in post-fidget response, we would expect to see distinct clusters of data points corresponding to each. Instead we find that when labeled by layer or area identity, the data was mixed and could not be visually separated---this implies that post-fidget responses did not differ between layer or area **(Supplementary Fig. 2)**. To validate that area or layer identity could not be distinguished based on post-fidget neuronal responses with UMAP, we trained a random forest classifier based on the UMAP-projected dataset using either the layer (175, 275, or 375 μm) or area as labels (VISp, VISpm, VISal, VISl). If differences between post-fidget responses for each of these layer/area classes exist, the model should achieve high classification accuracy. After training (see **Method**), we found instead that the classifier had low performance for both area and layer on test data, with comparable performance to the classifier trained on randomly permuted labels (**Fig 3e and 3f**).

Contrary to behaviors such as running, fidgets in passively viewing mice are non-relevant to vision. We therefore hypothesized that fidget neuronal response would not affect visual responses as opposed to running modulation. To do so, we compared the response of cells to different drifting grating orientations and spatial frequencies during baseline condition with during fidget (see **Fig 4a** for examples). We found that all 4 sub-types of fidget responsive cell were equally responsive to visual stimuli and their sensory tuning properties to drifting gratings (direction, orientation and temporal frequency selectivity) were not significantly different. (**Fig 4b-d**, p=0.24, 0.30, 0.06, 0,16, two-side t-test, comparing the response amplitude during fidget and baseline for active, phasic, depressed and neutral sub-class respectively). When comparing visual stimuli during and outside of fidget behaviors, the visually evoked responses were mostly identical (**Fig 4e)**. This is in contrast to a visually relevant behavior such as running, which significantly modulated visually evoked responses as reported previously^2,4^ (**Fig 4f**).

**Fig. 4.**
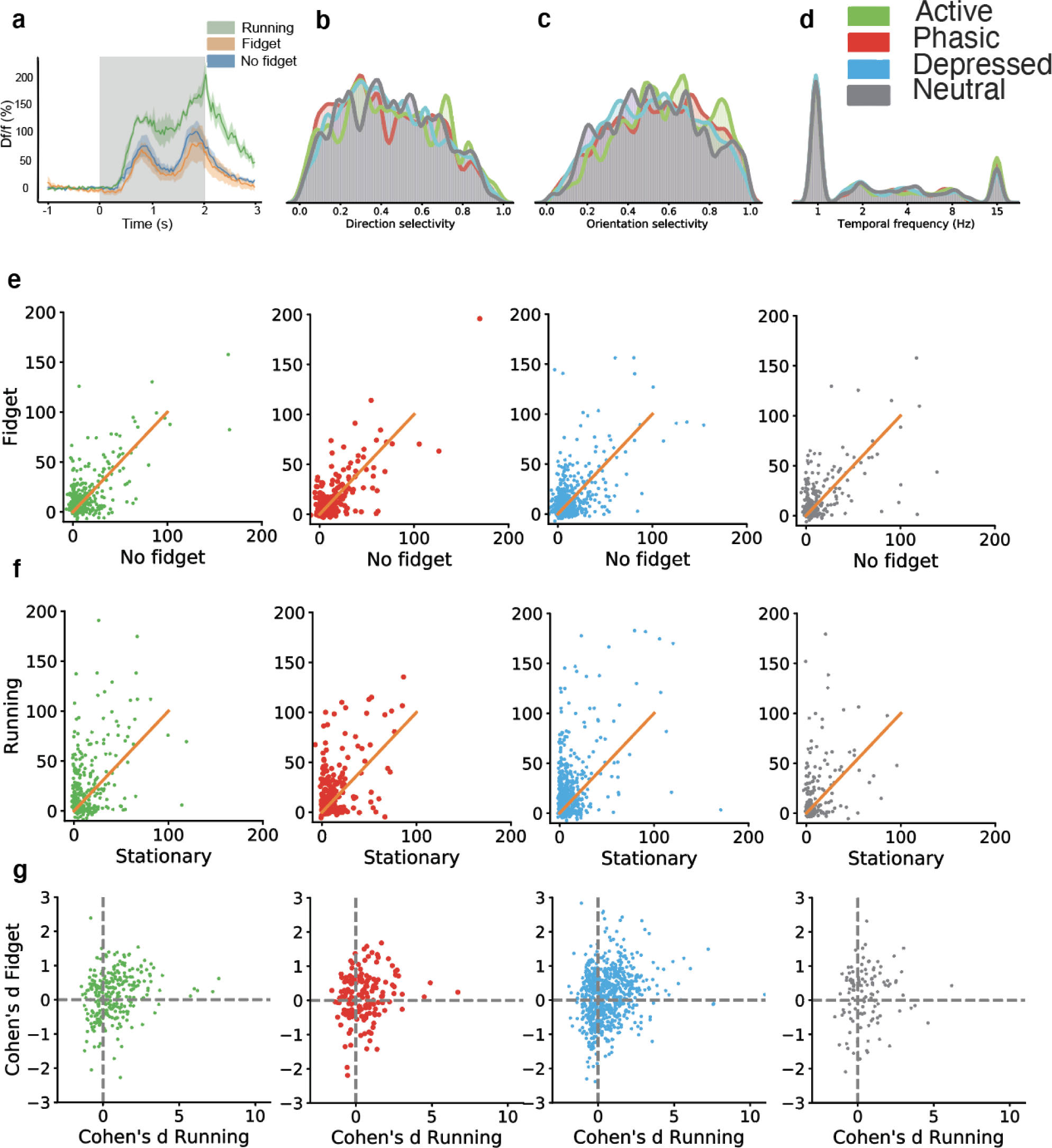
Contrary to running responses, neuronal fidget responses do not modulate visually evoked activity. **(a)** Example visual evoked responses of a cell to drifting gratings during running, fidgets, and resting conditions, shaded error bars indicate +/- SEM. **(b)** Normalized histograms of the direction selectivity of the four cells types during the drifting gratings stimulus is plotted **(c)** same as (b) but for cells’ preferred drifting grating orientation **(d)** same as (b) but for cells’ preferred temporal frequency of drifting gratings **(e)** The mean trial ΔF/F during fidget is plotted against non-fidget behavior for each cell during its preferred direction and temporal frequency of the drifting grating stimulus. The best fitting linear regression line is plotted in orange. This metric is plotted separately for all four cell types. **(f)** same as (d) but for running vs. stationary behaviors. **(g)** The Cohen’s d metric for fidget is plotted against the Cohen’s d for running behavior for each cell, across all four cell types.

Could a subset of the same cells be involved in behavior modulation in the visual cortex? To look at this, we compared the Cohen’s d of fidget (see **Methods**) and running evoked responses across the four identified neuron response types. Cells that were running modulated (p=0.0002, 0.0004, 0.13, 0,0009, two-side t-test, comparing the response amplitude during running vs stationary for active, phasic, depressed and neutral sub-class respectively) were in fact not modulated by fidget (p=0.24, 0.30, 0.06, 0.16, two-side t-test, comparing the response amplitude during running vs stationary for active, phasic, depressed and neutral sub-class respectively) (see **Fig 4g)**. In fact, among cells that were significantly modulated by running, only 4% of them were also significantly modulated by fidget.

In conclusion, our data supports that in passively viewing conditions, a behavioral event with no visual relevance broadly impacts the visual cortex but does not modulate visual responses.

## Discussion

Multiple recent studies have shown that motor activity greatly impacts the activity of not only the motor cortex but primary sensory areas like V1^2,8,30^. Given the rapid advance in brain-wide neuronal recording^21-22^, there has been increasing concern related to the importance of monitoring behavioral activity to account for spontaneous brain-wide neuronal activity^17^. Using a large two-photon Ca^2+^ imaging dataset collected in mice passively viewing a battery of standardized visual stimuli, we characterized the neuronal response of neurons of the visual cortex to fidgets, a single standardized motor output analogous to a startle response. We found that 47% of neurons show significant co-activity with fidgets throughout all areas and layers we recorded from. Previous studies in behaving mice have shown that brain-wide activity is better accounted for by uninstructed motor outputs than task driven signals^9^. Our study confirms the importance of taking into account motor activity when analyzing neuronal data.

We here propose a complementary approach to uncovering the role of motor signals in primary sensory areas. Many behavior tasks are associated with rich behavior outputs that can only be properly captured with multiple video cameras^8^. Even with appropriate monitoring, the dimensionality and variability of behavioral outputs make any interpretation more challenging. By focusing our analysis on a single standardized behavior event, we could group together a large number of behavioral trials, akin to visually-evoked stimulus trials. Closed-loop experimental designs could prove instrumental to extend this approach, by pairing specific visual stimuli with specific tracked behavioral events. Such an approach could allow a more direct analysis of how motor output modulates visual inputs.

“Visual” fidgets were previously analyzed in relation to visual stimuli^10^. They observed an increase of behavioral visual fidget response to gratings as supported here. Contrary to this study, all of our analysis were conducted in a familiar sensory environment where all mice were habituated to stimuli. We also included all fidgets in our subsequent analysis regardless of their timing with a visual stimulus. This allowed us to compare a large number of single cell responses to either fidgets or visual stimuli, extending this previous characterization to a novel direction.

We found that excitatory neurons were responding with 3 distinct temporal profiles to fidgets. Remarkably the proportion and responses of neurons in each class was maintained in all layers and brain areas we looked at, and consequently we could not predict the location of our recording using the response to fidget despite a large database to train our decoder on. This result suggests that behavioral information is not only broadcasted broadly, but also homogeneously throughout the cortical mantle. This result is to contrast with running modulation which typically impacts deeper layers more strongly^4^. Our interpretation is that local sensory inputs shape local behavioral representations depending on the causal overlap of those events. Future research could test whether a visually conditioned behavioral event transitions from a broad modulation to be more specific to certain visual areas during learning^24,25^. The broad availability of standardized behavior signals could be a key component in how the cortex selects, learns and binds local sensory information with relevant motor outputs.

## METHODS

### Transgenic mice

All animal procedures were approved by the Institutional Animal Care and Use Committee (IACUC) at the Allen Institute for Brain Science. Triple transgenic mice (Ai93, tTA, Cre) were generated by first crossing Ai93 mice with Camk2a-tTA mice, which preferentially express tTA in forebrain excitatory neurons. Double transgenic mice were then crossed with a Cre driver line to generate mice in which GCaMP6f expression is induced in the specific populations of neurons that express both Cre and tTA.

Rorb-IRES2-Cre;Cam2a-tTA;Ai93 (n=10) exhibit GCaMP6f in excitatory neurons in cortical layer 4 (dense patches) and layers 5,6 (sparse). Cux2-CreERT2;Camk2a-tTA;Ai93 (n=16) expression is regulated by the tamoxifen-inducible Cux2 promoter, induction of which results in Cre-mediated expression of GCaMP6f predominantly in superficial cortical layers 2, 3 and 4. Slc17a7-IRES2-Cre;Camk2a-tTA;Ai93 (n=2) is a pan-excitatory line and shows expression throughout all cortical layers. Scnn1a-Tg3-Cre;Camk2a-tTA;Ai93 (n=5) exhibit GCaMP6f in excitatory neurons in cortical layer 4 and in restricted areas within the cortex, in particular primary sensory cortices. Nr5a1-Cre;Camk2a-tTA;Ai93 (n=1) exhibit GCaMP6f in excitatory neurons in cortical layer 4. Rbp4-Cre;Camk2a-tTA;Ai93 (n=11) exhibit GCaMP6f in excitatory neurons in cortical layer 5. Ntsr1-Cre_GN220;Ai148 (n=1) exhibit CaMP6f in excitatory corticothalamic neurons in cortical layer 6.

### Animal head-implants and cortical window implantation

Transgenic mice expressing GCaMP6f were weaned and genotyped at ~p21, and surgery was performed between p37 and p63. Surgical protocols were described in previous publications associated with the two-photon datasets^4^.

### Intrinsic imaging and mapping of the visual cortex

Retinotopic mapping was used to delineate functionally defined visual area boundaries and enable targeting of the in vivo two-photon calcium imaging to retinotopically defined locations in primary and secondary visual areas. Retinotopic mapping protocols were described in previous publications associated with the two-photon datasets^4^.

### In vivo two-photon chronic imaging

Calcium imaging was performed using a two-photon-imaging instrument, Nikon A1R MP+. The Nikon system was adapted to provide space to accommodate the behavior apparatus). Laser excitation was provided by a Ti:Sapphire laser (ChameleonVision – Coherent) at 910 nm. Pre-compensation was set at ~10,000 fs2. Movies were recorded at 30Hz using resonant scanners over a 400 μm field of view.

Mice were head-fixed on top of a rotating disk and free to walk at will. The disk was covered with a layer of removable foam (Super-Resilient Foam, 86375K242, McMaster) to alleviate motion-induced artifacts during imaging sessions.

An experiment container consisted of three imaging sessions (60 min each) at a given field of view during which mice passively observed three different stimuli. The same location was targeted for imaging on all three recording days to allow repeat comparison of the same neurons across sessions. One imaging session was performed per day, for a maximum of 16 sessions for each mouse.

On the first day of imaging at a new field of view, the ISI targeting map was used to select spatial coordinates. A comparison of surface vasculature patterns was used to verify the appropriate location by imaging over a field of view of ~800 μm using epi-fluorescence and blue light illumination. Once a cortical region was selected, the imaging objective was shrouded from stray light from the stimulus screen using opaque black tape. In two-photon imaging mode, the desired depth of imaging was set to record from a specific cortical depth. On subsequent imaging days, we returned to the same location by matching (1) the pattern of vessels in epi-fluorescence with (2) the pattern of vessels in two photon imaging and (3) the pattern of cellular labelling in two photon imaging at the previously recorded location.

Calcium imaging data was collected at the four cortical depths of 175, 275, 350 and 375 micrometers. Throughout our analysis, data from the cortical depth of 175 micrometers were classified as layer 2/3, 275 and 350 micrometers as layer 4, and 375 as layer 5.

More details on our imaging protocols are available in our previous publication^4^.

### Visual stimulation

Visual stimuli were generated using custom scripts written in PsychoPy as described previously^4^.

Visual stimuli included drifting gratings, static gratings, locally sparse noise, natural scenes and natural movies. These stimuli were distributed across three ~60 minutes imaging sessions. During session A the drifting gratings, natural movie one and natural movie three stimuli were presented. During session B the static gratings, natural scenes, and natural movie were presented. During session C the locally sparse noise, natural movie one and natural move two were presented. In each session, the different stimuli were presented in segments of 5-13 minutes and interleaved with each other. In addition, at least 5 minutes of spontaneous activity were recorded in each session.

#### Drifting Gratings

The total stimulus duration was 31.5 minutes. The stimulus consisted of a full field drifting sinusoidal grating at a single spatial frequency (0.04 cycles/degree) and contrast (80%). The grating was presented at 8 different directions (separated by 45°) and at 5 temporal frequencies (1, 2, 4, 8, 15 Hz). Each grating was presented for 2 seconds, followed by 1 second of mean luminance gray before the next grating. Each grating condition (direction & temporal frequency combination) was presented 15 times, in a random order. There were blank sweeps (i.e. mean luminance gray instead of grating) presented approximately once every 20 gratings. This stimulus was used to measure the direction tuning, orientation tuning and temporal frequency tuning of the cells.

#### Static Gratings

The total stimulus duration was 26 minutes. The stimulus consisted of a full field static sinusoidal grating at a single contrast (80%). The grating was presented at 6 different orientations (separated by 30°), 5 spatial frequencies (0.02, 0.04, 0.08, 0.16, 0.32 cycles/degree), and 4 phases (0, 0.25, 0.5, 0.75). The grating was presented for 0.25 seconds, with no inter-grating gray period. Each grating condition (orientation, spatial frequency, and phase) was presented ~50 times in a random order. There were blank sweeps (i.e. mean luminance gray instead of grating) presented roughly once every 25 gratings. This stimulus was used to measure the spatial frequency tuning and the orientation tuning of the cells, providing a finer measurement of orientation than provided from the drifting grating stimulus.

#### Locally Sparse Noise

Stimulus. The total stimulus duration was 37.5 minutes. The Locally Sparse Noise stimulus consisted of a 16 × 28 array of pixels, each 4.65 degrees on a side. For each frame of the stimulus (which was presented for 0.25 seconds), a small number of pixels were white, a small number were black, and the rest were mean gray. The white and black spots were distributed such that no two spots were within 5 pixels of each other.

#### Natural Scenes

The stimulus consisted of 118 natural images. Images 1-58 were from the Berkeley Segmentation Dataset, images 59-101 from the van Hateren Natural Image Dataset, and images 102-118 are from the McGill Calibrated Colour Image Database. The images were presented in grayscale and were contrast normalized and resized to 1174 × 918 pixels. The images were presented for 0.25 seconds each, with no inter-image gray period. Each image was presented ~50 times, in random order, and there were blank sweeps (i.e. mean luminance gray instead of an image) roughly once every 100 images.

#### Natural Movie

Three different clips were used from the opening scene of the movie Touch of Evil (Welles, 1958). Natural Movie 1 and Natural Movie 2 were both 30 second clips while Natural Movie 3 was a 120 second clip. All clips had been contrast-normalized and were presented in grayscale at 30 fps. Each movie was presented 10 times in a row with no inter-trial gray period.

### Behavioral monitoring

During calcium imaging experiments, eye movements and animal posture were recorded. The left side of each mouse was imaged with the stimulation screen in the background to provide a detailed record of the animal response to all stimuli. The eye facing the stimulus monitor (right eye) was recorded using a custom IR imaging system. No pupillary reflex was evoked by any of these illumination LEDs.

Eye tracking video hardware includes a camera (Allied Vision, Mako G-032B with GigE interface) acquiring at a rate of 30 fps, with a 33ms exposure time and gain between 10-20. These videos were illuminated using an LED (Engin Inc, LZ1-10R602) at a wavelength of 850 nm with a fixed lens in front (Thorlabs, LB-1092-B-ML).

Animal behavior monitoring hardware includes a camera (Allied Vision Mako G-032B with GigE interface) with a 785 nm short pass filter (Semrock, BSP01-785R-25), and lens (Thorlabs MVL8M23, 8mm EFL, f/1.4). The camera acquires at a rate of 30 fps, with a 33ms exposure time and a set gain of 10. The short-pass filter is affixed to the camera to suppress any light from the eye tracking LED. Illumination for the behavior monitoring camera comes from a 740 nm LED (LED Engine Inc, LZ4, 40R308-000). A bandpass filter (747 +/- 33nm, Thorlabs, LB1092-B-ML) is affixed in front of the illumination LED to prevent visible portion of the LED spectrum from reaching the mouse eye.

Two-photon movies (512×512 pixels, 30Hz), eye tracking (30 Hz), and a side-view full body camera (30 Hz) were recorded and continuously monitored.

### Processing of two-photon calcium imaging movies

For each two-photon imaging session, the image processing pipeline performed: (1) spatial or temporal calibration, (2) motion correction, (3) image normalization to minimize confounding random variations between sessions, (4) segmentation of connected shapes and (5) classification of soma-like shapes from remaining clutter.

The motion correction algorithm relied on phase correlation and only corrected for rigid translational errors. Each movie was partitioned into 400 consecutive frame blocks, representing 13.3 s of video. Each block was registered iteratively to its own average three times. A second stage of registration integrated the periodic average frames themselves into a single global average frame through six additional iterations. The global average frame served as the reference image for the final resampling of every raw frame in the video.

Fluorescence movies were processed using a segmentation algorithm to identify somatic regions of interest (ROIs) that was described previously^2^. Segmented ROIs were matched across imaging sessions. For each ROI, events were detected from ΔF/F by using an L0-regularized algorithm. For each neuron, we z-score ΔF/F trial activity and compute the mean z-scored response of each neuron aligned to the time of fidget onset (0 seconds).

To determine the significance of neural activity modulation post-fidget, we apply several threshold criteria to clustered neuronal activity. The threshold criteria were used in previous calcium imaging literature, and here we present two that gave very different results: One criteria where the mean ΔF/F is larger than 6%, and one criteria where the maximum ΔF/F during the post-fidget period is greater than 5%.

### Behavioral Analysis

We use histogram of oriented gradients (HOG) descriptors as our model features due to their invariance to position, rotation, scale, and changes to lighting between mice. Additionally, HOG vectors have been shown to perform well in dynamic behavioral classification across different individuals^4^.

Side-view full-body camera (30 Hz) videos were converted to grayscale, normalized using power law compression before processing, and manually cropped to exclude background elements including the screen and rotation disk. Cropping was done by a manual selection tool that constructed a rectangle from four clicked points. Pixels outside this rectangle were cropped out. The four points were systematically chosen in this order: upper-left as the eye of the mouse, lower-left as the closest point on the running disk forming a line to the first chosen point that is parallel to the vertical axis, lower-right as the point forming a line to the second point that is parallel to the surface of the running disk, and finally upper-right as the closest point intersecting the head-stage forming a line to the third point that is parallel to the vertical axis. Histograms of Oriented Gradients (HOGs) were then computed for each frame using the following parameters: 8 histogram orientation bins, using square cells with a height of 32 pixels and 1 cell per block, and then dimensionality reduced using principal component analysis (50 dimensions). The features were then concatenated in one second blocks and fed into the model.

The training set of ~ 100,000 video frame labels was collected from six human annotators who used a custom XML based tool. A radial basis function support vector machine (SVM) was trained on the training set with a cross-entropy loss function (C and gamma parameters of the SVM found using a grid search).

The normalized fidget rate was computed by subtracting the baseline fidget rate during inter-stimulus grey stimulus presentation and dividing by the standard deviation of the fidget rate.

To estimate fidget magnitude, an optical flow measure was computed for each grayscale cropped frame using python’s OpenCV Optical flow function. We used the Gunner’s Farneback algorithm using Two-Frame Motion Estimation Based on Polynomial Expansion with a 30 pixels kernel size. For each example, the optical flow measure was integrated over the duration of all continuous frames labeled as fidget.

### Fidget neuronal response analysis

Neural responses were aligned to the onset of the fidget behavior and cropped to keep 100 frames (~ 3 seconds) preceding the initiation of the fidget and 200 frames (~ 6 seconds) post fidget initiation.

Neural activity was normalized on a trial-by-trial basis by subtracting the mean activity of the 100 frames (~3 seconds) of baseline neural activity preceding the initiation of the fidget response and dividing by the standard deviation of activity. Across trial z-scored neural activity was then averaged to get the mean z-scored activity for each neuron.

### Clustering of fidget neuronal responses

The mean z-scored activity for all neurons post-fidget was passed into the k-means++ clustering algorithm with clustering evaluated using the gap statistic. The optimal number of clusters was found to be four.

### Analysis of visually evoked responses in the visual cortex along with their fidget and running modulation

To compare visually evoked responses during fidget and non-fidget, we computed the mean ΔF/F trial activity during frames annotated as fidget and non-fidget. Specifically, we only compute this metric during cells’ preferred direction and temporal frequency tuning for the drifting gratings stimulus, defined as the stimulus parameters which elicit the largest mean response.

To investigate whether there is an interaction between fidget and running modulation, we computed a Fidget Cohen’s d metric defined as:

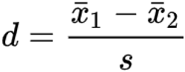

where *<u>x</u>*<u>1</u> is the mean ΔF/F during fidget, *<u>x</u>*<u>2</u>is the mean ΔF/F during non-fidget and *s* is the norm of the standard deviation of ΔF/F during fidget and non-fidget. We then computed the running Cohen’s d metric same as above but for running vs. stationary behavior. For each cell, we then plotted the fidget and running Cohen’s d against each other per individual cell, and computed a covariance that indicates whether cell’s that were highly modulated by fidget were also highly modulated by running.

### UMAP and Random Forest Classification

To search for any differences between fidget neural responses across different cortical layers and visual areas we took 200 frames (~ 6 seconds) of each neuron’s normalized average post fidget activity and passed them into a nonlinear dimensionality reduction method, UMAP. In Python, we used the umap package (umap 0.4.0rc3, hyperparameters min_dist = 0.0 and n_neighbors = 200) to project the dataset into a 2-dimensional embedding to elicit any differences in the data. The embedded data points were then passed into a random forest classifier. An 85-15 train-test split was used along with 5-fold cross-validation with XGBoost (xgboost 1.2.0) and Scikit-Learn (sklearn 0.23.2). A grid hyperparameter search was used to optimize the classifier. To counter class-imbalance in the training and test sets, each class was randomly subsampled down to the count of the least prevalent class in the set. After training, F1-scores (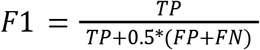; TP, FP, and FN are true positives, false positives, and false negatives respectively) were estimated for each class.

### Statistical tests

Multiple comparisons were corrected using the Benjamini–Hochberg false discovery rate framework (*q* < 0.05), and all statistical tests in the study were two-tailed, two-sample Kolmogorov–Smirnov tests.

## Data availability

The behavioral data that support the findings of this study are available from the corresponding authors upon request.

## Acknowledgements

We thank the Allen Institute founder, Paul G. Allen, for his vision, encouragement, and support. We thank Fuhui Long for support and guidance in training SVM. We thank Eric Shea-Brown and Adrienne Fairhall for continuous support and encouragement. We thank Amy Bernard for initial internship support. We thank Doug Ollerenshaw for comments on the manuscript. We thank Ahmed Al Dulaimy for his coding expertise on the video annotator.

## Author Contributions

M.R., J.L. designed and conceptualized analyses. M.R. designed fidget annotation. M.R analyzed all fidget and neuronal data. E.K.L. performed cluster validation analysis. S.C., I.K., P.R., K.R., F.G, T.V.N, J. Larkin., K.M., A.K., R.H., N.B. annotated behavioral data. M.R., J.L., E.K.L, S.C. wrote the paper. J.L. supervised the research.

