## supplement for "A standardized non-visual behavioral event is broadcasted homogeneously across cortical visual areas without modulating visual responses"

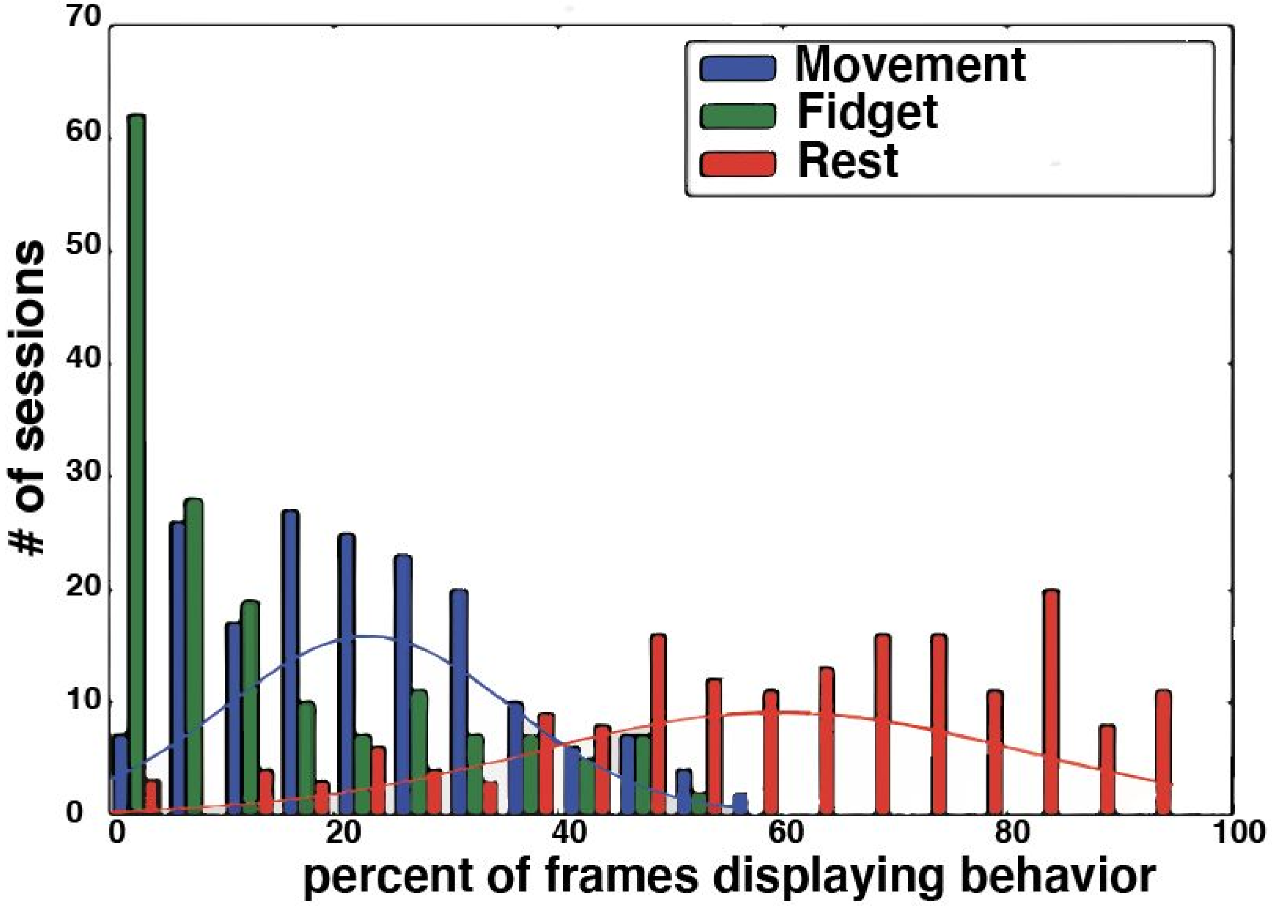


**Supp. Fig. 1 | Fidget rate frequency across sessions. (a)** (left) Bar plot visualizing the number of sessions (total of 144) with their percentage of video frames classified as one of three behaviors (movement, resting, or fidget).

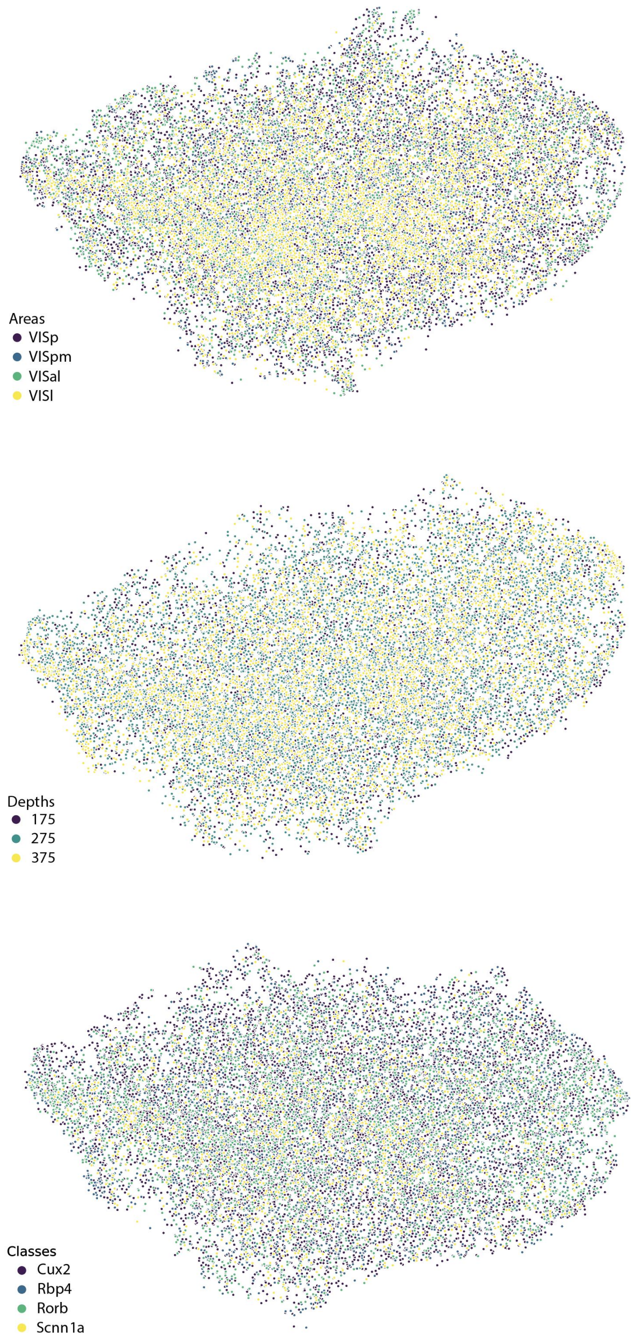


**Supp. Fig. 2 | UMAP embeddings of post-fidget single neuron responses by area, layer, and mouse Cre-line.**

Visualization of the projection of post-fidget neural responses from averaged single neurons into a 2D embedded space identified by unsupervised non-linear dimensionality reduction (UMAP), labeled by area (top subplot), layer (middle subplot), and mouse Cre-line (bottom subplot).

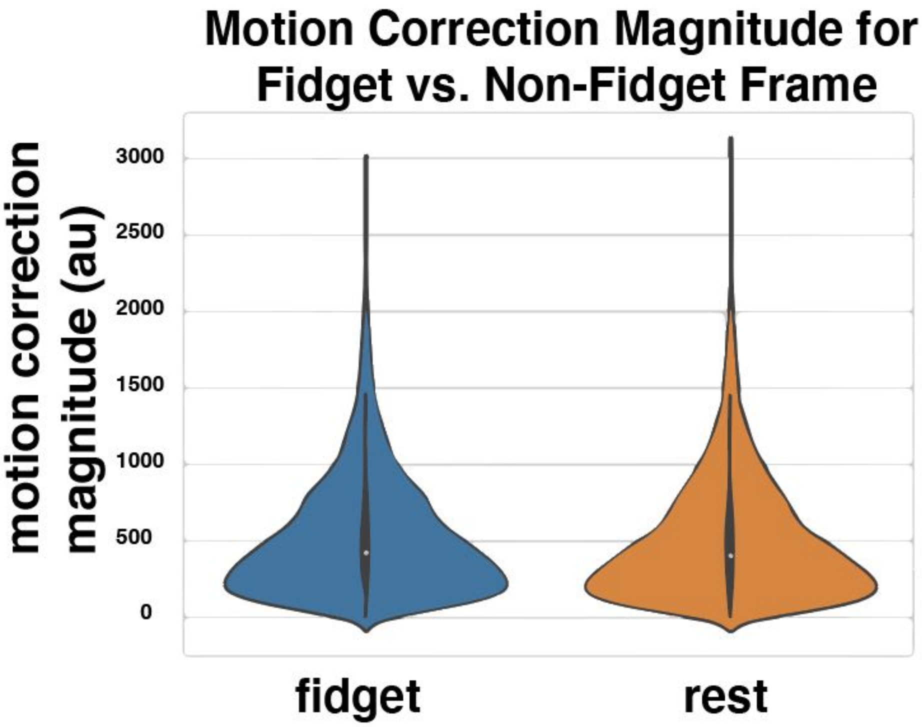


**Supp. Fig. 3 | Fidgets do not induce translational movement of the cortex.**

Violin plots of the Euclidean distance of 2P image motion in the x and y dimensions (translational movements) during mouse fidget vs. mouse resting state (no movement).

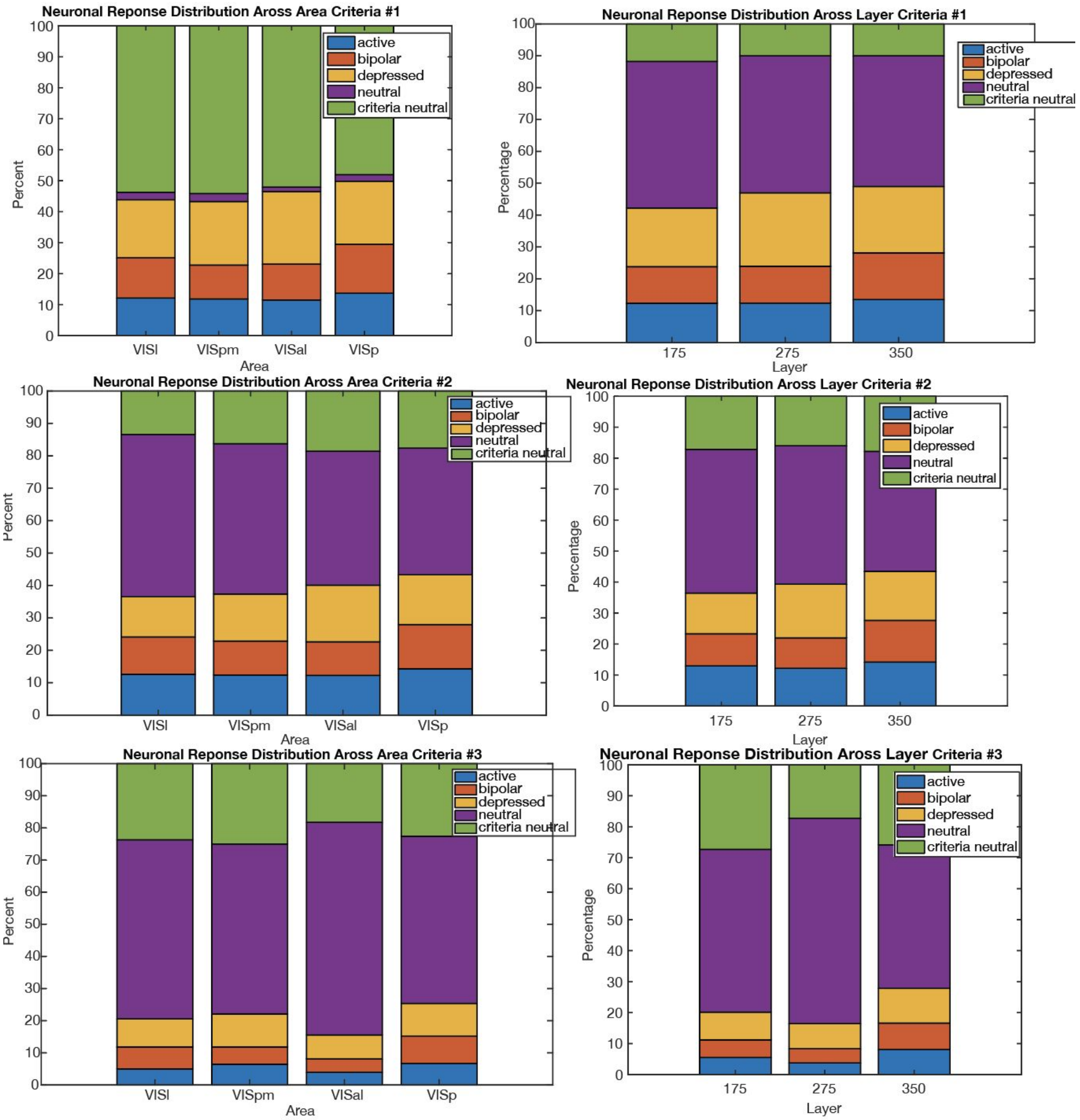


**Supp. Fig. 4 | Equal distribution of clustered neuronal response types across area and layer is robust to threshold criteria.**

Distributions of clustered neuronal responses after applying three different threshold criteria to neural data. The distributions are conditioned in area or layer, and neural responses that did not meet the threshold criteria are labeled in green as criteria neutral. Depending on the strictness of the criteria the percent of criteria neutral neurons changes, but the distribution of neural response types conditioned on area or layer remains fairly consistent.

https://www.dropbox.com/sh/1o6ph7vnxlgk4vl/AAChvNj2BBq9dtv4tbZxpWU6a?dl=0

**Supp. video. 1 | Example live calcium imaging of neuronal activity during fidget (fidget initiation indicated).**

The size of each calcium imaging frame is 400 μm x 400 μm. Videos are trial-averaged calcium responses during fidget (-3 (s) to +6 (s) relative to fidget initialization) across an imaging session. Videos frames are converted to greyscale, played at 2x speed and compressed as a .mp4 file.
